# Listening shapes seeing: Sustained auditory spatial attention enhances early visual-cortical processing

**DOI:** 10.64898/2026.08.11.744194

**Authors:** Yong Min Choi, Viola S. Störmer

## Abstract

How does the auditory system implement spatial selection without a dedicated cortical map for space? One hypothesis holds that auditory spatial attention draws on a supra-modal network including the parietal-occipital cortex; an alternative implicates subcortical structures (e.g., superior colliculus) with no direct recruitment of visual cortex. To adjudicate between these accounts, we used a dichotic listening paradigm and tested whether sustained auditory spatial attention produces the behavioral and neural signatures of visual spatial attention, which would only be expected if auditory attention engages the same cortical mechanisms. Participants listened to two digit streams, spoken by male and female voices, played from left and right speakers. They were instructed to attend to either the left stream, the right stream, or a specific voice gender. A behavioral experiment (N=24) showed higher discrimination accuracy for visual stimuli appearing intermittently at the auditorily attended relative to unattended location. Furthermore, participants’ gaze was reliably biased towards the attended location. In a second experiment (N=14), we used electrophysiological recordings of frequency-tagged visual evoked potentials to more directly assess early visual processing, and found enhanced visual-cortical responses for stimuli matching the location of the attended auditory stream. In addition, occipital alpha power (8–11 Hz) was reduced over the hemisphere contralateral to the attended sound stream. Together, these effects mirror the hallmarks of visual-spatial attention, suggesting that auditory spatial attention co-opts the architecture of the visual cortex to implement spatial selection, thereby directly enhancing visual processing.

**Significance statement:** Human can effortlessly direct spatial attention to a sound’s location in the external world. Yet the auditory system has no dedicated spatial map in the brain, raising a fundamental question: how does auditory spatial attention arise? We show that sustained attention to a sound based on its location produces well-known signatures of visual spatial attention: enhanced visual-perceptual sensitivity, larger early visual-cortical responses, and modulation of occipital alpha-band activity and oculomotor behavior. This converging behavioral and neural evidence demonstrates that auditory spatial attention actively engages and reshapes early visual processing, pointing to a supra-modal attention system shared across the senses.

## Introduction

In everyday environments, sensory inputs from multiple sources compete for limited cognitive resources, requiring the brain to selectively prioritize relevant inputs over irrelevant ones (Carrasco, 2011; Desimone & Duncan, 1995). One key mechanism for achieving this is spatial selection, where information is prioritized based on its location in space (Posner, 1980). In vision, such spatial selection can be guided by the retinotopic organization of visual cortex, which provides a near one-to-one mapping between the locations in external space and internal neural representations (DeYoe et al., 1996; Martinez et al., 1999; Sereno et al., 1995; Wandell et al., 2007). Audition, by contrast, lacks an analogous spatial map. Although binaural cues (i.e., interaural time or intensity differences) provide a computational basis for inferring sound location, no topographic cortical map of auditory space has been identified (Kong et al., 2014; Middlebrooks, 2021; Middlebrooks et al., 1998). This poses a fundamental puzzle for cognitive neuroscience: how does the auditory system implement spatial selection when, unlike vision, it lacks a dedicated cortical map for space?

One hypothesis holds that auditory spatial attention draws on a supra-modal attention network that shares a common neural architecture with visual spatial selection (Banerjee et al., 2011; Kong et al., 2014; Lee et al., 2014; Popov et al., 2023; Shomstein & Yantis, 2004; Smith et al., 2010). Consistent with this view, parietal cortex appears to play a central role in multi-sensory spatial attention: parietal lesions impair spatial orienting of attention in both the visual and auditory modality (Farah et al., 1989), parietal TMS can causally disrupt auditory spatial attention (Deng et al., 2019), and even in congenitally blind individuals, auditory spatial attention engages the frontal eye fields and medial occipital cortex (Garg et al., 2007) – regions that form part of the visual-spatial attention network (Corbetta & Shulman, 2002). Other work has shown that auditory spatial attention can modulate visual cortex for peripheral sounds outside the visual field of view (Cate et al., 2009). Together, these findings suggest that parietal and occipital regions – brain areas that are retinotopically organized – may play a role not just in visual but also auditory spatial attention.

An alternative hypothesis suggests that auditory spatial attention primarily involves subcortical structures such as the superior colliculus (SC; Basso & May, 2017; Krauzlis et al., 2013; Rinne et al., 2008) or its non-mammalian homolog, the optic tectum (Knudsen & Brainard, 1991), which contain well-defined spatial maps and are known to be involved in spatial attention across modalities. Acoustic cues are first integrated into a map of auditory space, which is then relayed to the deep layers of the SC, where it becomes aligned with topographically organized visual and somatosensory maps (Knudsen, 2018). Furthermore, not only multisensory input converges at subcortical structures, the SC also receives descending signals from forebrain areas conveying spatial behavioral goals, and may therefore support spatial selection in multisensory contexts specifically (Knudsen, 2018; Stein et al., 1993). On this view, auditory spatial attention would not necessarily modulate brain areas traditionally associated with visual-spatial attention, such as early visual cortex, a prediction our study directly tests.

A hallmark of visual spatial attention is its direct and spatially selective influences on visual-perceptual processing (Anton-Erxleben & Carrasco, 2013; Carrasco, 2018). Visually attending to a location in space enhances both behavioral performance and early visual-cortical responses to objects appearing at the attended location. Additionally, maintaining spatial attention on either visual hemifield is known to be accompanied by biased fixational oculomotor behavior (Awh et al., 2006; Moore et al., 2003) and by a reduction in occipital alpha-band activity over the hemisphere contralateral to the attended location (Worden et al., 2000). A particularly strong test of whether auditory spatial attention shares these neural mechanisms is therefore to examine whether it produces comparable behavioral and neural consequences in early visual cortex. If it does, this would provide compelling evidence that auditory-spatial attention is implemented, at least in part, through the same retinotopically organized visual cortical architecture that underlies visual spatial attention.

To test this, we employed a dichotic listening task in which participants performed a digit 1-back task while attending to a voice stream either based on its location (left, right) or non-spatial feature (gender: female, male). Visual processing was probed in two complementary experiments: psychophysically, using an orientation discrimination task alongside measures of oculomotor behavior, and neurally by examining steady-state visual evoked potentials (SSVEPs) as a marker of early visual-cortical processing (Andersen et al. 2011), and occipital alpha-band oscillations, an established signature of visual-spatial attention (Worden et al., 2000).

Our results provide converging evidence that sustained auditory spatial attention recruits visual structures in a spatially specific manner, thereby modulating early visual processing. Visual discrimination accuracy was higher and eye gaze was reliably biased towards the attended auditory stream, though these gaze biases did not explain the visual performance benefits. SSVEP amplitudes were larger for visual stimuli appearing in the hemifield congruent with the attended auditory stream. In addition, auditory spatial attention reduced the occipital alpha-band rhythm contralateral to the attended location — the canonical neural signature of visual spatial attention. Together, our findings support the view of a supra-modal spatial attention system in which sustained auditory selection recruits the cortical architecture traditionally known to support visual-spatial attention. Critically, this shared recruitment does not only involve higher-order parietal spatial maps, but extends to early visual cortex, directly shaping visual perception itself.

## Results

### Experiment 1: Behavioral study

Twenty-four healthy adults completed a behavioral experiment designed to measure the visual-perceptual consequences of sustained auditory spatial attention. Two concurrent streams of digits were presented through speakers located on the left and right side of the monitor (Figure 1A) and participants performed an auditory 1-back task, detecting immediate digit repetitions while maintaining attention on a single digit stream. Before each block, they were instructed to attend to the auditory stream from (1) the left speaker (attend-left), (2) the right speaker (attend-right), or (3) to the stream spoken by a specific voice (male or female; attend-voice). For example, in the attend-left block, subjects attended to the voice coming from the left speaker and pressed the spacebar whenever the same digit was repeated twice in a row regardless of voice type, while ignoring digits coming from the right speaker. The attend-voice block served as a non-spatial control as male/female voices were randomly intermixed across both speakers within each block, keeping sensory input identical between blocks.

**Figure 1.**
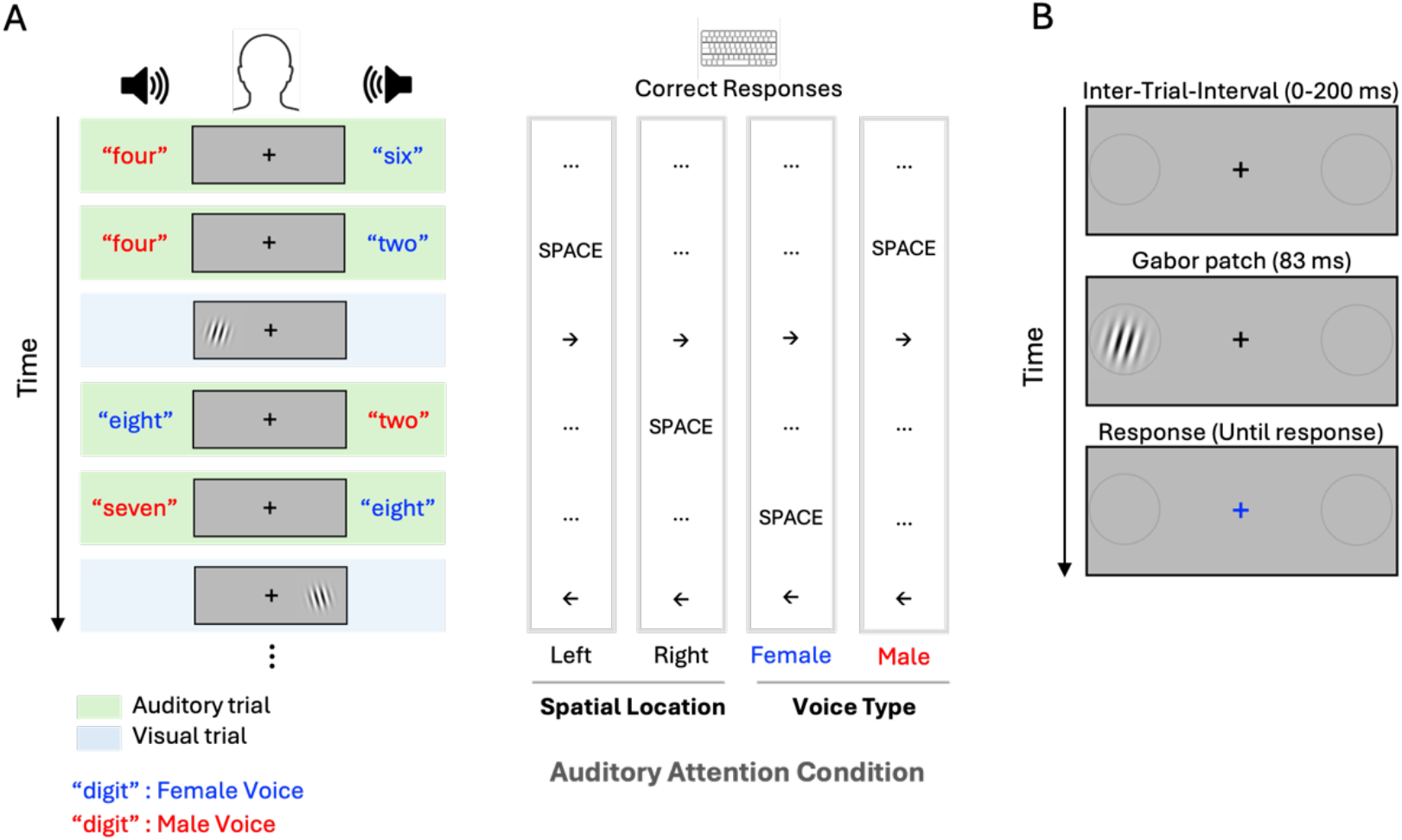
Behavioral experiment design. (A) Each block consisted of intermixed auditory and visual task trials. Throughout the block, two concurrent streams of spoken digits — one in a male voice and one in a female voice — were presented from speakers positioned to the left and right of the monitor. Before each block, participants were instructed to attend to one of three auditory streams: the left speaker, the right speaker, or a specific voice gender (male or female) regardless of location. Participants pressed a key whenever the same digit repeated consecutively within the attended stream (auditory 1-back task). (B) On interleaved visual trials, a sub-threshold Gabor patch was briefly flashed in either the left or right visual hemifield, and participants reported its orientation.

Performance on the auditory 1-back task was well above chance, with a high hit rate (M = 0.70, SD = 0.03) and a low false alarm rate (M = 0.03, SD = 0.02). Auditory task sensitivity (d-prime) did not significantly differ between auditory attention conditions (F(2, 46) = 0.794, p = .458, *η_p_^2^* = 0.033), indicating comparable task difficulty across attention conditions and no systematic performance differences attributable to the locus of auditory spatial attention.

### Sustained auditory spatial attention modulates visual discrimination performance

Concurrent with the auditory task, visual sensitivity was assessed using an intermittent orientation discrimination task (Figure 1B). On each visual trial, a near-threshold Gabor patch was briefly presented in either left or right visual hemifield, and participants reported its orientation (clockwise or counterclockwise). Visual trials were randomly interleaved among auditory trials (52 visual and 156 auditory trials per block), ensuring that the auditory task remained the primary focus throughout.

We computed orientation discrimination accuracy for each Gabor location and auditory attention condition (Figure 2A). Then, for the statistical evaluation, discrimination accuracies were collapsed across left and right locations and attention conditions, yielding three conditions: valid (auditory and visual locations *matched*), invalid (auditory and visual locations *mismatched*), and control (attend-voice) (Figure 2B). Performance in the attend-voice blocks served as a baseline to assess whether observed effects reflected facilitation or suppression. A repeated-measure ANOVA revealed a significant main effect of attention condition on visual discrimination accuracy (F(2, 46) = 4.288, p = .020, *η_p_^2^* = 0.157). Post hoc comparisons showed that accuracy in the valid condition was significantly higher than invalid (t(23) = 3.355, p_bonf_ = .008, d = 0.202) and the control condition (t(23) = 2.598, p_bonf_ = .048, d = 0.169). No significant difference was observed between the invalid and control conditions (t(23) = −0.349, p_bonf_ = 1.000, d = −0.032). These results provide behavioral evidence suggesting enhanced visual processing at spatial locations that match the locus of sustained auditory spatial attention.

**Figure 2.**
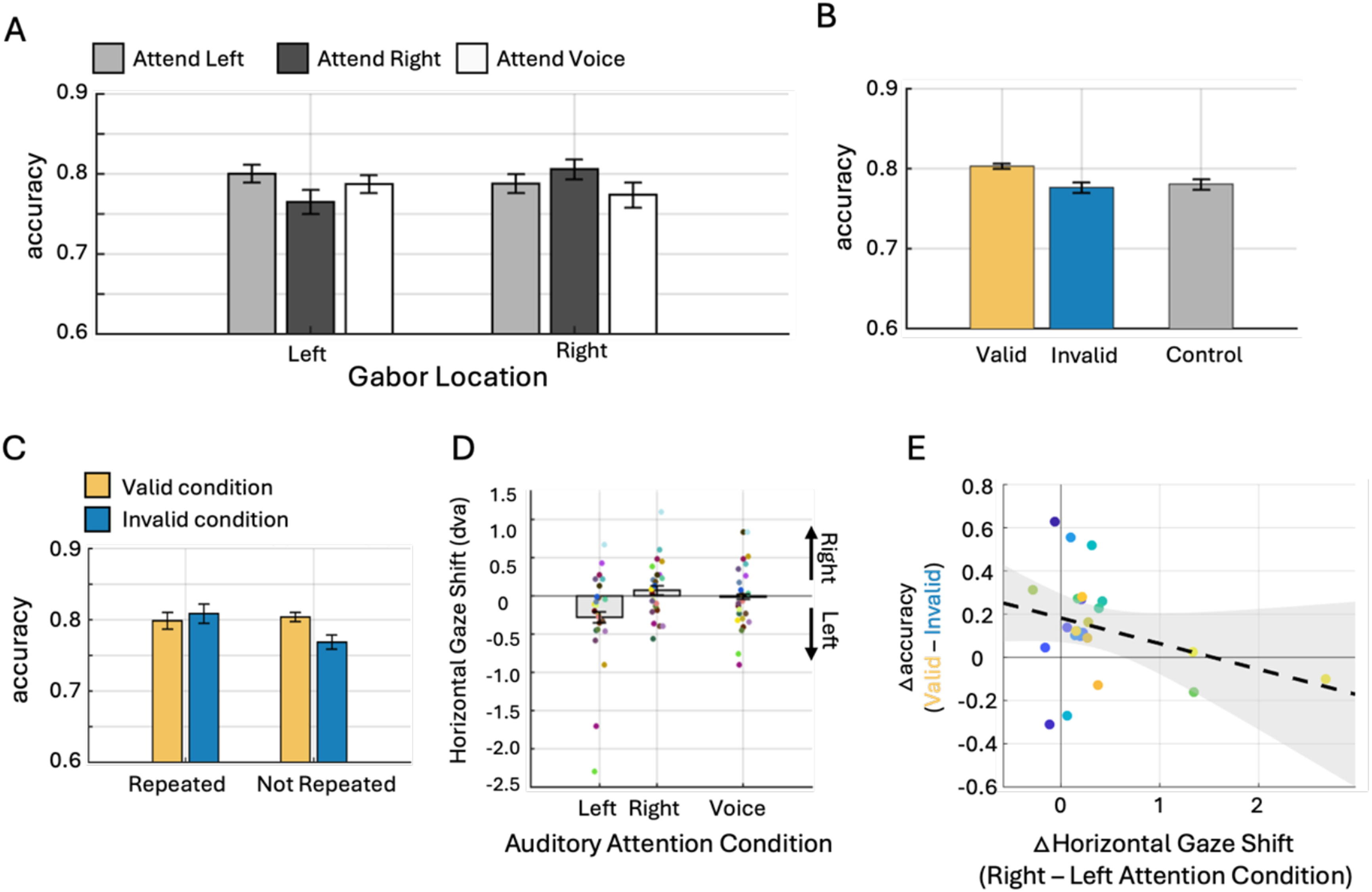
Behavioral experiment results (N=24). (A) Orientation discrimination accuracy across visual target location and auditory attention conditions. (B) Trials were grouped as valid (Gabor patch was at attended location) or invalid (Gabor patch elsewhere); attend-voice trials served as baseline. (C) Trials were split based on whether the previous trial involved a repeated auditory target (repeated) or not (not repeated). (D) Horizontal gaze shifts during the pre-stimulus window (−500 to 0 ms) toward left (negative) or right (positive) visual field as a function of auditory attention condition. (E) Scatter plot showing the relationship between gaze shifts (x-axis) and the effect of sustained auditory attention on visual discrimination (y-axis). Dots represent participants; error bars indicate within-subject standard error.

### Sustained, rather than temporary shift of attention modulates visual discrimination performance

A possible alternative to a sustained-attention account is that visual performance was instead boosted by temporary shifts of auditory attention (Spence & Driver, 1997; McDonald et al., 2013; Störmer et al., 2009). The dichotic listening paradigm presented constant auditory sensory inputs while focusing on the effect driven by the location of endogenous spatial attention. Nevertheless, it is still possible that the occurrence of digit repetition in the auditory stream (the target event) transiently shifted spatial attention toward the attended location and modulated visual processing of the subsequent visual trial.

To test this, we sorted visual trials according to the nature of the immediately preceding auditory event: repeated (the 1-back target) or non-repeated (Figure 2C). If a temporary transient attentional shift following the presentation of task-relevant target drives subsequent visual benefit, the effect of auditory spatial attention (valid vs invalid) should be larger when immediately followed by repeated auditory trials than by non-repeated trials. A 2×2 repeated-measures ANOVA with attention validity (valid and invalid) and preceding-trial type (repeated and non-repeated) as within-subject factors yielded a significant interaction (F(1, 23) = 4.312, p = .049, *η_p_^2^* = 0.158). Interestingly, post-hoc comparisons showed a significant effect of auditory spatial attention only in non-repeated trials (t(23) = 4.106, p_bonf_ = .003, d = 0.258), but not in repeated trials (t(23) = −0.498, p_bonf_ = 1.000, d = −0.073).

Note that because only 20% of auditory digits were repeated, the repeated condition contained far fewer trials (M = 61.9, SD = 5.9) than the non-repeated condition (M = 250.1, SD = 5.9). To account for this imbalance, we iteratively subsampled the non-repeated trials to match the number of repeated trials and found results consistent with the first analysis (Supplementary Figure 1). This bootstrapped result confirms that cross-modal facilitation cannot be attributed to brief temporary shifts of attention, and supports the conclusion that sustained, goal-directed auditory spatial attention systematically enhances visual processing at the attended location.

### Gaze shift toward auditory spatial attention

Fixational oculomotor behavior (i.e., gaze shift) is another hallmark of visual spatial attention (Awh et al., 2006; Bollimunta et al., 2018; Krauzlis et al., 2013; Moore et al., 2003). Thus, if auditory spatial attention recruits a common neural substrate with visual spatial attention, an analogous gaze shift toward the spatial locus of auditory attention would be expected. To test this, we examined gaze position during the 500-ms time window^1^ preceding stimulus onset and found a small yet reliable leftward gaze shift when attending to the left auditory stream relative to the right (t(23) = −2.786, p = .011, d = −0.569; Figure 2D).

Given this systematic gaze shift, we next asked whether it underlies the enhanced visual discrimination performance: attention-driven gaze shifts could, in principle, bring the visual probe closer to fixation (i.e., reduce its retinal eccentricity) and thereby improve discrimination accuracy in the valid condition. Although this account seems unlikely given how small the gaze shift was (attend-right minus attend-left: M = 0.35, SD = 0.62 dva), we nonetheless tested this directly by asking whether the magnitude of the gaze shift correlated with the size of auditory-attention effects on visual discrimination performance. A scatter plot (Figure 2E) suggested no positive relationship; if anything, the trend was slightly negative (*r* = −0.35, *p* = 0.089, BF_10_ = 1.88). Thus, although auditory spatial attention reliably produced small gaze shifts at fixation, this oculomotor response did not appear to account for the observed perceptual enhancement, suggesting that the two effects are dissociable and may reflect independent signatures of visual-spatial attention mechanisms engaged by auditory spatial attention.

### Experiment 2: EEG study

The behavioral data showed that visual discrimination performance was enhanced in the hemifield that was spatially congruent with the attended auditory stream, suggesting that auditory spatial attention has direct influences on visual behavior. This behavioral finding raises several important questions. Is the cross-modal facilitation contingent on the visual stimulus being task-relevant? Does this facilitation arise at the level of early visual-cortical processing, or does it, at least in part, reflect post-perceptual processes such as decision-making (Matsumiya & Furukawa, 2023), motor planning, or response execution (Pashler, 1994; Richardson et al., 2013)? Does sustained auditory spatial attention modulate the same neural processes in occipital cortex that have been identified for visual-spatial attention (Worden et al., 2000; Thut et al., 2006)?

To answer these questions, fourteen healthy adults completed a similar dichotic listening task while EEG activity was recorded. To probe the implicit neural consequences of sustained auditory spatial attention on visual processing, two task-irrelevant checkerboards were continuously presented in the left and right visual hemifield (Figure 3A). The two checkerboards flickered at distinct frequencies (12 Hz or 15 Hz; Figure 3B), eliciting separable steady-state visual evoked potentials (SSVEPs). At the beginning of every 10-second trial, an instruction text indicated whether participants should attend to the left (attend-left) or right (attend-right) auditory stream for the auditory 1-back task while maintaining their gaze at the central fixation cross.

**Figure 3.**
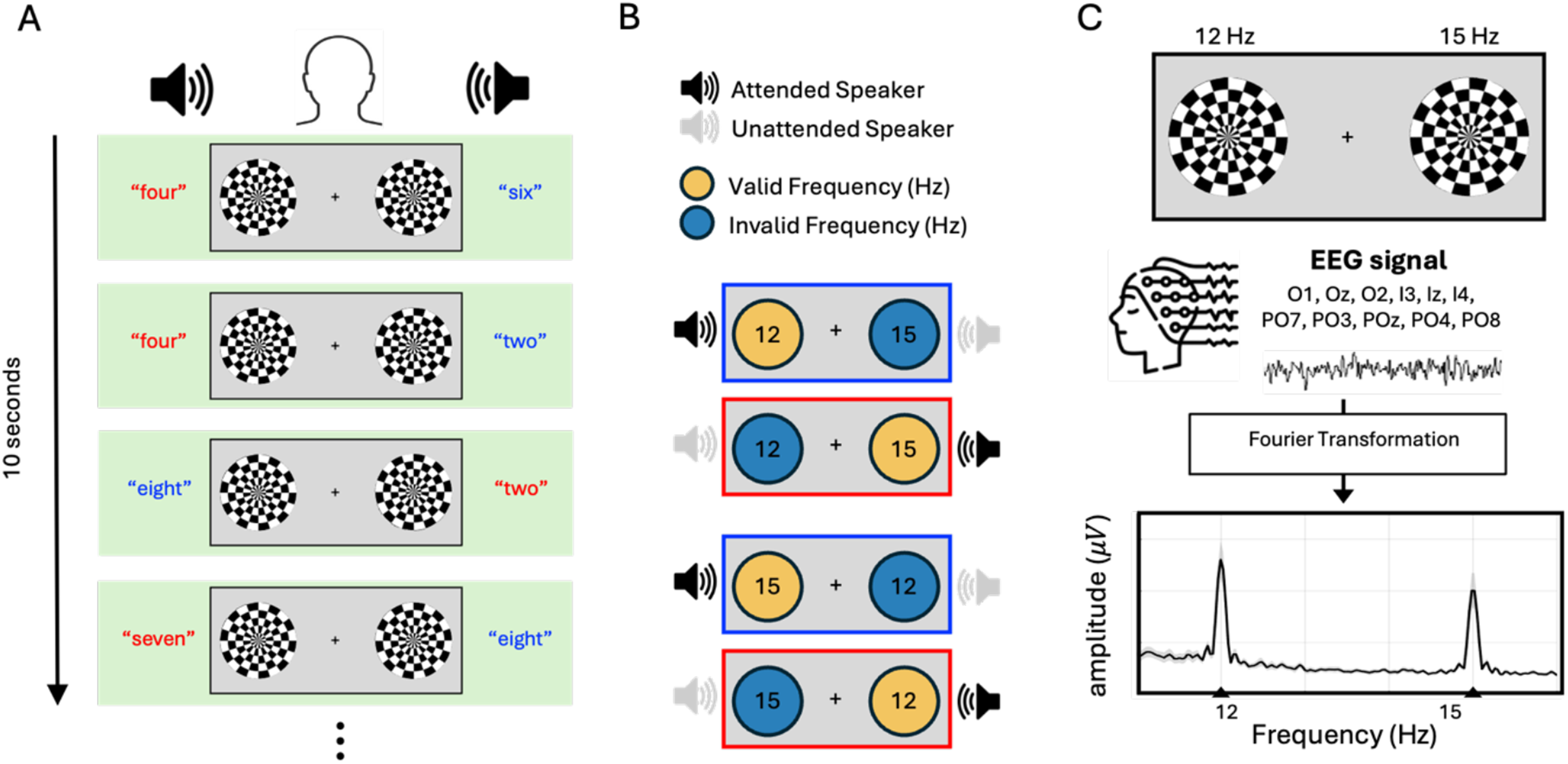
EEG experiment design. (A) Auditory 1-back task with task-irrelevant visual stimulation. Each trial consisted of 10 s of an auditory 1-back task while participants passively viewed checkerboard patterns flashing in the left and right visual hemifields. (B) Attention manipulation and frequency tagging. Participants attended to either the left or right auditory stream, while the two checkerboards flickered at 12 Hz and 15 Hz. Frequencies were labeled as valid or invalid based on the correspondence between the attended auditory location and the spatial location of each flicker. (C) Schematic of Steady-State Visual Evoked Potential (SSVEP) analysis. EEG signals from each trial and electrode were Fourier-transformed to obtain amplitude spectra. The bottom panel shows amplitude spectra pooled across occipito-parietal electrodes and conditions, with the black line indicating the grand average across participants, revealing peaks at 12 Hz and 15 Hz. Peak amplitude reflects the strength of visual processing in each hemifield.

### Sustained auditory spatial attention enhances SSVEP amplitude in the congruent hemifield

To determine whether the locus of auditory attention modulates early visual processing in each hemifield, we analyzed the amplitude of SSVEPs at the tagged frequencies (12 Hz and 15 Hz). EEG data were averaged across trials associated with one of the four conditions (Figure 3B), and Fourier-transformed to obtain amplitude spectra (Figure 3C). Because the checkerboard stimuli flickered at 12 and 15 Hz, the spectra exhibited corresponding peaks at those frequencies, with peak amplitude indexing the strength of early visual encoding for each hemifield (Figure 3C, bottom panel).

Figure 4A shows amplitude spectra around the tagged frequencies for both flicker assignments (12/15 Hz and 15/12 Hz across hemifields) over occipito-parietal electrodes (O1, Oz, O2, I3, Iz, I4, PO7, PO3, POz, PO4, PO8). In both assignments, SSVEP amplitudes were larger in the hemifield spatially congruent with the attended auditory stream. For example, when the left stimulus flickered at 12 Hz (upper panel), the 12 Hz amplitude was greater for attend-left than attend-right trials, and this pattern remained consistent regardless of tagged frequencies and their assigned hemifield. Topographical maps of SSVEP amplitude further confirmed that responses were maximal over occipito-parietal regions (Figure 4B).

**Figure 4.**
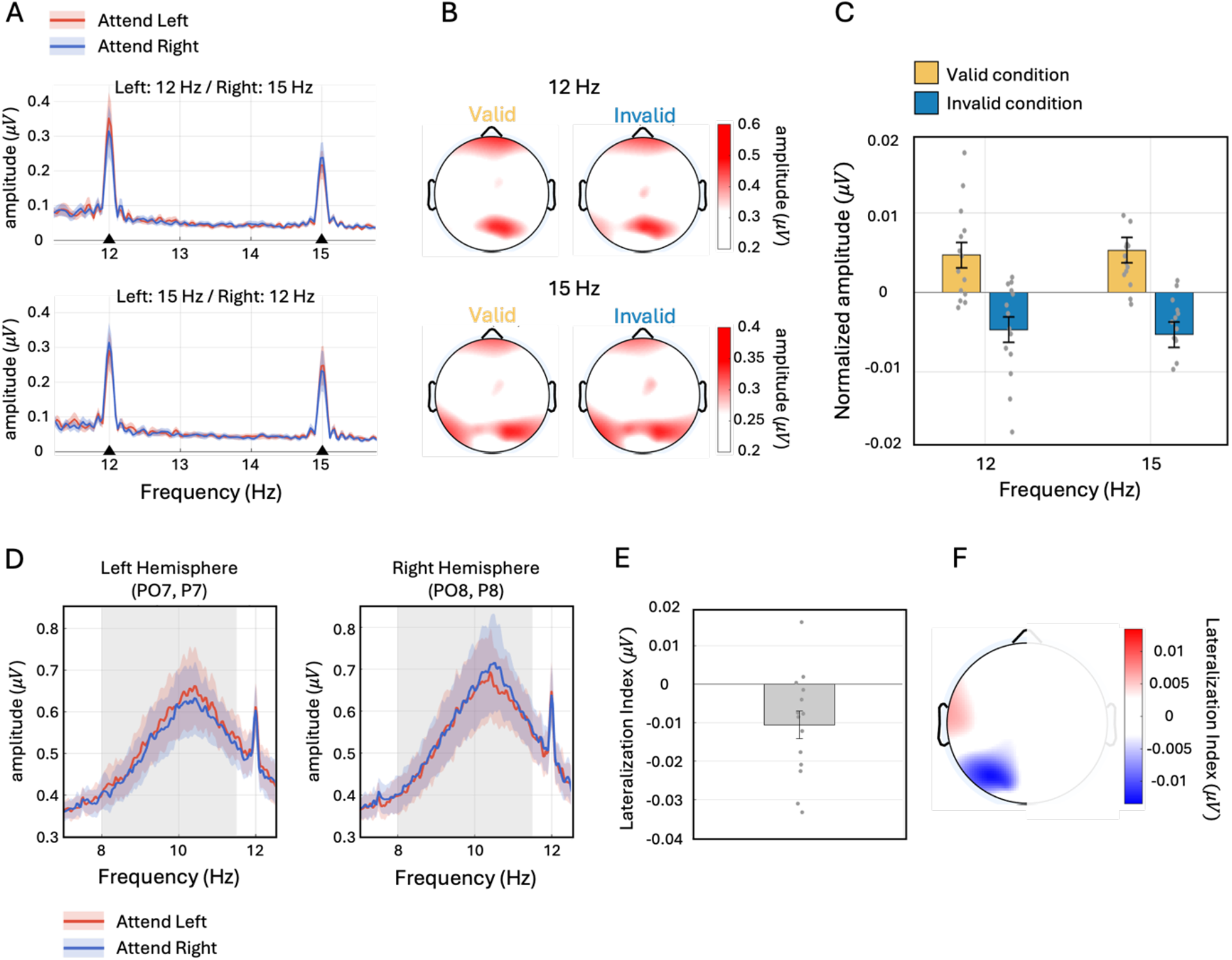
EEG experiment results (N = 14). (A) Amplitude spectra around the tagged frequencies (12 Hz and 15 Hz). The upper panel shows trials in which the left and right checkerboards flickered at 12 Hz and 15 Hz, respectively; the lower panel shows the reverse frequency assignment. Within each panel, blue and red lines indicate trials in which participants attended to the left or right speaker, respectively. (B) Topographical maps of SSVEP amplitude. The upper row shows 12 Hz amplitude for valid (left) and invalid (right) trials; the lower row shows the corresponding 15 Hz maps. (C) Normalized SSVEP amplitude. Amplitudes at each tagged frequency were normalized and grouped into valid and invalid conditions according to the spatial correspondence between the attended location and stimulus location. (D) Alpha-band (8–11.5 Hz) amplitude spectra averaged across occipito-parietal electrodes over the left (PO7, P7) and right (PO8, P8) hemispheres (Keefe & Störmer, 2021). Blue and red lines indicate attention directed to the left and right speaker, respectively. (E) Alpha lateralization index, computed as the difference between alpha-band amplitude contralateral and ipsilateral to the attended hemifield, divided by their sum. (F) Topographic scalp map showing the spatial distribution of alpha lateralization, collapsed across left and right electrode pairs, and projected to the left side of the scalp. In all bar plots, individual dots represent single participants and error bars denote within-subject standard error; shaded regions in amplitude spectra denote ±1 standard error.

For statistical comparison, we normalized SSVEP amplitudes separately for each participant and tagged frequency (12 Hz and 15 Hz). To normalize across the two conditions which differed only in the locus of auditory spatial attention, we first computed the mean SSVEP amplitude across the attended and unattended conditions. We then subtracted this mean from the amplitude in each condition and divided the result by the same mean, yielding a normalized (percent-change-like) measure for each condition (cf., Andersen et al., 2011; Störmer et al., 2014). Normalized amplitudes were then averaged across occipital-parietal electrodes and collapsed into valid and invalid conditions (Figure 4C). A 2×2 repeated-measures ANOVA revealed a significant main effect of attention condition (F(1, 13) = 12.69, p = .003, *η_p_^2^* = 0.494, BF_incl_ = 10.71), with no significant attention condition × frequency interaction (F(1, 13) = 0.146, p = .708, *η_p_^2^* = 0.011, BF_incl_ = 0.429)^2^. The same pattern of results held when analyzing each electrode group (O, I, and PO) separately (Supplementary Figure 3).

Greater SSVEP amplitudes for stimuli in the visual hemifield congruent with the attended auditory location indicate that sustained auditory spatial attention enhances visual-cortical processing in the congruent hemifield, even when the visual modality is completely task-irrelevant. Furthermore, the result points to a genuine modulation of early visual-cortical processing, similar to what has been observed in visual-spatial attention paradigms.

### Auditory spatial attention reduces contralateral posterior alpha-band activity

Next, we examined posterior alpha-band activity over electrodes contralateral vs. ipsilateral with respect to the attended auditory location (Feng et al., 2017; Keefe et al, 2021; Störmer et al., 2016), a well-established neural signature of the allocation of visual spatial attention (Worden et al., 2000). Unlike the SSVEP analysis, Fourier transforms were applied to single-trial EEG data, as endogenous alpha oscillations are not phase-locked between trials. We focused on alpha frequencies between 8-11.5 Hz, avoiding contamination from visual-tagged frequency (12 Hz), at parietal–occipital electrodes (PO7, PO8, P7, P8; Keefe & Störmer, 2021).

Amplitude spectra averaged across trials revealed a reduced alpha-band amplitude over electrodes located contralateral to the attended auditory location (Figure 4D). That is, alpha-band amplitude was attenuated over left-hemisphere electrodes when attention was directed to the right relative to the left auditory stream (Figure 4D, left), and vice versa. To quantify these effects, we averaged alpha amplitudes (8–11.5 Hz) across all electrodes and computed a lateralization index (LI) as the difference between attending to the contralateral vs. ipsilateral hemifield. Negative LI values indicate reduced alpha-band amplitude over the posterior cortex contralateral to the attended auditory stream. We found a reliable lateralization of alpha-band activity as a function of auditory attention condition: the LI was significantly negative over parietal-occipital electrodes (PO7/PO8/P7/P8; t(13) = −2.831, p = .014, d = −0.757, BF_10_ = 4.278), indicating that alpha-band amplitude was systematically reduced over the occipital cortex contralateral to the attended auditory stream. The topographical scalp maps further confirmed the modulation of alpha power over posterior electrode sites, indicating that the observed lateralized alpha activity was primarily generated by neural sources in occipital-parietal cortex (Figure 4E). We found no lateralized alpha amplitude from electrodes over auditory cortex (T7,T8; t(13) = 1.724, p = .108, d = 0.461, BF_10_ = 0.881).

The reduction of posterior alpha power contralateral to the attended auditory location mirrors the canonical neural signature of visual spatial attention (Worden et al., 2000; Thut et al., 2006). Thus, maintaining auditory attention at a specific spatial location does not only biases behavioral performance and enhance visual-cortical processing, as indexed by the SSVEP amplitude changers, but also engages the same alpha-based inhibitory mechanism that the visual system uses to prioritize relevant space. This convergence provides strong evidence that sustained auditory spatial attention invokes neural mechanisms that are functionally equivalent to those underlying visual spatial attention.

## Discussion

How does the auditory system implement spatial selection when, unlike vision, it lacks a dedicated cortical map for space? One possibility is that auditory spatial attention draws on subcortical structures such as the superior colliculus, which contains spatial maps formed by converging signals from multiple sensory systems (Basso & May, 2017; Krauzlis et al., 2013). Alternatively, auditory spatial attention may recruit the same cortical regions as visual-spatial attention, modulating neural activity in parietal and occipital areas – regions traditionally associated with visual-spatial attention. (Banerjee et al., 2011; Farah et al., 1989; Kong et al., 2014; Lee et al., 2014; Popov et al., 2023; Shomstein & Yantis, 2004). In the current study, we directly tested this by examining whether auditory spatial attention gives rise to the known behavioral and neural hallmarks of visual spatial attention.

### Auditory spatial attention enhances early visual processing at the congruent location

Experiment 1 demonstrated that attending auditorily to a specific location has direct consequences for visual perception: Visual discrimination performance was significantly higher when a near-threshold Gabor patch appeared in the hemifield matching the attended auditory stream (valid) compared to both the mismatched hemifield (invalid) and the non-spatial control condition (attend-voice). Critically, the invalid condition did not differ from the control, indicating *facilitation* at the attended location rather than suppression at the unattended location.

To identify the neural signature of visual facilitation during an auditory attention task, Experiment 2 measured visual-cortical processing directly using electrophysiological recordings. SSVEP amplitudes were larger for visual stimuli presented in the hemifield congruent relative to incongruent with the attended auditory location. This result demonstrates that sustained auditory spatial attention modulates early visual-cortical processing itself, rather than post-perceptual stages such as decision-making (Matsumiya & Furukawa, 2023) or motor preparation (Pashler, 1994). Moreover, this modulation occurred even though the visual stimuli were entirely task-irrelevant, suggesting that auditory spatial attention automatically enhances visual-cortical responses in a spatially specific manner. As changes in SSVEP amplitude are a well-established index of gain modulations in early visual cortex due to feature and spatial attention in the visual domain (Morgan et al., 1996; Andersen et al., 2011; Störmer et al., 2013; Özkan & Störmer 2024; Özkan, Chapman & Störmer, 2025), the observed increases in SSVEP amplitudes imply that auditory spatial attention can drive similar gain changes in early visual cortex as visual attention. Furthermore, this indicates that auditory spatial attention does not only recruit higher-order parietal spatial maps, as previously shown, but also extend to earlier stages of visual processing that directly shape visual perception itself.

### Auditory spatial attention modulates alpha-band activity over visual cortex

We also found reduced alpha-band activity (8–11.5 Hz) over occipito-parietal cortex contralateral to the attended auditory stream. Lateralized alpha activity is a well-established marker of endogenous visual spatial attention, often interpreted as a preparatory gain signal that biases competition between attended and unattended locations (Desimone & Duncan, 1995; Jensen & Mazaheri, 2010; Kastner et al., 1999; Kelly et al., 2006; Thut et al., 2006; Worden et al., 2000).

This alpha-lateralization finding also connects to previous literature linking auditory spatial attention to lateralized alpha activity. Auditory spatial attention has been shown to produce parietal alpha lateralization that shifts systematically with the attended location (Banerjee et al., 2011; Mehraei et al., 2018; Deng et al., 2020; Popov et al., 2023), and disrupting parietal alpha with transcranial magnetic stimulation causally impairs auditory spatial attention (Deng et al., 2019). Our finding that this lateralized alpha modulation extends to occipital cortex suggests that oscillatory dynamics traditionally attributed to visual spatial selection also support spatial attention in the auditory domain. Furthermore, combined with the behavioral facilitation and early visuo-cortical modulation reported above, the current study demonstrates that auditory spatial attention does not merely exploit spatial maps in parietal cortex, but also brings about measurable changes over visual regions themselves.

### Sustained and reflexive effects of auditory spatial attention on visual processing

The present result relates to prior research showing that brief, task-irrelevant sounds that involuntarily capture spatial attention improve visual detection and discrimination at the same location (Frassinetti et al., 2002; Feng et al., 2014; McDonald et al., 2000; Spence & Driver, 1997; Störmer et al., 2009). Similar to the current study, the behavioral benefits of exogenous auditory cueing were also accompanied by increased visual-cortical responses over the hemisphere contralateral to the sound’s location (Feng et al., 2014; Keefe et al., 2021; McDonald et al., 2013; Störmer et al., 2009) as well as reduced occipital alpha-band power (8–14 Hz) contralateral to the attended hemifield (Feng et al., 2017; Keefe et al., 2021; Störmer et al., 2016).

Yet, these previous studies focused entirely on involuntary, transient shifts of spatial attention, leaving open the question of whether voluntary, sustained auditory attention produces comparable effects. Real-world listening often demands sustained spatial attention to a specific location — for example, when focusing on a target voice among competing speakers (Bronkhorst, 2000; Cherry, 1953). Given that exogenous and endogenous spatial attention rely on partially distinct neural mechanisms (Corbetta & Shulman, 2002; Chica et al., 2013; Bowling et al., 2020; Tang et al., 2016), and behavioral dissociations between the two have been reported across multiple studies (see Table 1 in Chica et al., 2013), it remained important to understand whether sustained auditory spatial attention engages the same visual cortical mechanisms as its involuntary counterpart.

**Table 1.**
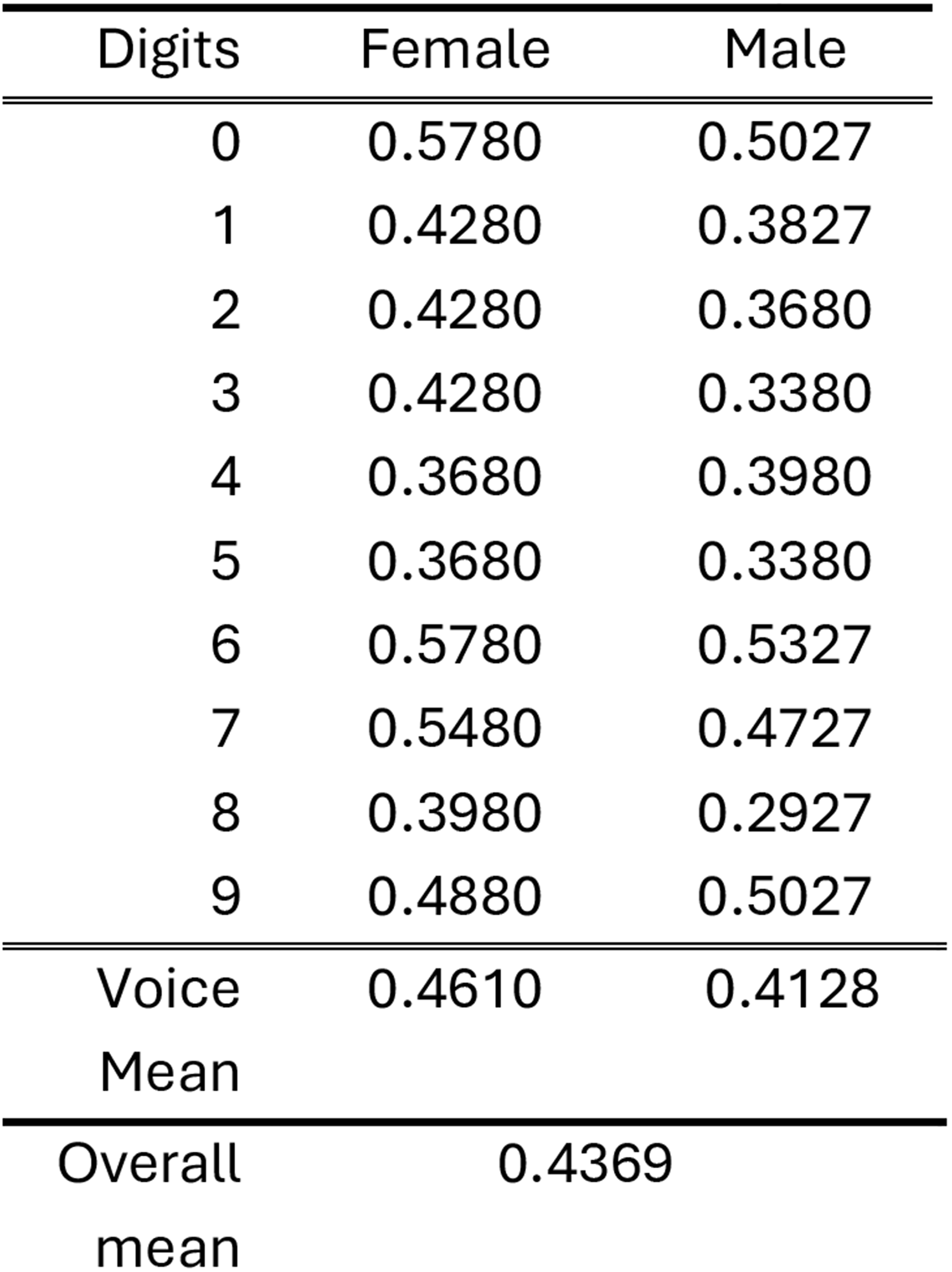
Auditory stimulus duration (sec) after preprocessing.

Using a dichotic listening paradigm, we directly tested this and found converging evidence that sustained auditory spatial attention enhances visual processing in a spatially specific manner. Together with prior work on exogenous auditory cueing then, our results support a view in which auditory spatial attention — whether allocated voluntarily and in a sustained manner or involuntarily and reflexively — activates visual-cortical processes similar to those engaged by visual spatial attention.

### Gaze shift tracks auditory spatial attention

Simultaneous eye-tracking data revealed gaze shifts toward the spatial hemifield attended auditorily, consistent with prior literature (Popov et al., 2023). Systematic gaze shifts represent another signature of visual spatial attention engagement (Awh et al., 2006; Bollimunta et al., 2018; Krauzlis et al., 2013; Moore et al., 2003). Deploying covert spatial attention in the visual modality biases the direction of fixational eye movements, not only when attention is directed to the external world (Corneil & Munoz, 2014; Hafed et al., 2011; Engbert & Kliegl, 2003), but also when it is directed to internal representations maintained within the spatial layout of working memory (van Ede et al., 2019; van Ede et al., 2020; van Ede et al., 2021). Critically, a recent primate study found that covert spatial attention enhanced visuo-cortical responses when accompanied by a microsaccade toward the attended location, providing strong support for an obligatory link between oculomotor bias and the neural signatures of visual spatial attention (Lowet et al., 2018); however, other work suggests that although fixational eye movements correlate with attention-related alpha-band activity, they are not strictly necessary for such modulation to occur (Liu et al., 2022; Liu et al., 2024). Even on this more conservative view, the correlational link between the brain’s oculomotor and visual-spatial attention systems raises the possibility that auditory spatial attention recruits the same oculomotor-linked visual system that underlies visual spatial attention. In our data, the observed gaze shifts were relatively small — within the fixational range (< 0.5 dva) — and their magnitude was not correlated with the visual facilitation effect. This suggests that the oculomotor bias does not account for the observed perceptual enhancement, but rather that the behavioral and oculomotor results may be dissociable and thus reflect independent signatures of visual-spatial attention mechanisms recruited by auditory spatial attention.

### Visual-cortical recruitment supports a supra-modal attention system

The current study was motivated by the logic that if subcortical structures were the primary substrate of the supra-modal spatial attention system, auditory spatial attention would not necessarily modulate cortical areas traditionally associated with visual-spatial attention. Our findings provide direct evidence for the recruitment of occipito-parietal cortex. The most parsimonious explanation for why auditory task demands alone modulate visual processing — in the absence of any visual relevance — is that a shared spatial attention mechanism spans across sensory modalities. Meanwhile, we should be careful not to overinterpret these findings. The design and results of the current study cannot rule out the possibility that subcortical structures also play a critical role in mediating spatial attention across the senses (Lomber et al., 2001). Indeed, the SC projects signals to visual cortex via the pulvinar nucleus of the thalamus, raising the possibility that the cortical recruitment we observed is itself inherited from subcortical structures routing signals into cortex through this ascending tecto-pulvinar pathway (Hu et al., 2019; Krauzlis et al., 2013). Regardless, our results provide clear evidence that auditory spatial attention affects visual cortex activity and, as a consequence, alters visual perception.

### Functional significance of cross-modal facilitation

What would be the functional role of cross-modal facilitation mediated by a supra-modal spatial attention system? We speculate that the effects we observed reflect an ecologically adaptive coupling between the senses. In the current experiment setting, visually attending to the congruent hemifield conferred no explicit task advantage. However, in natural environments, sensory inputs from the same spatial location typically belong to the same event, and selectively enhancing visual processing at the attended auditory location would facilitate multisensory integration (Mahoney et al., 2021). Such audio-visual coupling may also underlie other well-known cross-modal perceptual phenomena in which co-localized inputs complement one another: the perceived location of a sound is biased toward a concurrent visual stimulus (Hairston et al., 2003; Rohe & Noppeney, 2015), lip movements transform auditory perception as in the McGurk effect (McGurk & MacDonald, 1976), and sounds bias the interpretation of ambiguous visual stimuli (Williams et al., 2022, 2024). By enhancing sensory processing at the spatial focus of attention regardless of modality, the system may be optimized to exploit these cross-modal interactions whenever they arise.

## Conclusion

Our results are clear: where we listen shapes how we see. Voluntarily sustaining auditory attention at a specific location – for example when listening to someone in a noisy, crowded room – enhances visual-cortical processing and visual performance at the corresponding location. Most importantly, the effects on visual processing mirror the neural consequences typically observed in visual spatial attention tasks, suggesting that auditory spatial attention engages the same cortical mechanisms traditionally attributed to the visual domain. Together, our results provide direct support for a supra-modal spatial attention system shared across the senses, implemented at least in part through the recruitment of occipito-parietal cortex.

## Materials and Methods

### RESOURCE AVAILABILITY

All experimental materials, including MATLAB scripts for stimulus presentation, raw data, and analysis code are made publically available on the Open Science Framework (OSF): [https://osf.io/r2uhk/overview?view_only=f359a75f07994466b717c099cd109bc9].

### SUBJECT DETAILS

All participants were recruited from Dartmouth College, and reported to have normal or corrected-to-normal visual acuity. We collected informed consent in accordance with the procedures approved by the Institutional Review Board at Dartmouth College and were compensated with either course credit or monetary rewards for their participation.

Twenty-four participants (21 women and 3 men; *M*_age_ = 18.62 years, *SD*_age_ = 0.77) completed the behavioral experiment. The required sample size was calculated before data collection based on the effect of endogenous auditory spatial attention on visual discrimination performance (t(15) = 3.45, p = .004, d’ = 0.86; Keefe & Störmer, 2021), using G*Power version 3.1.9.6 (Faul et al., 2009). A priori power analysis, assuming a reduced effect size in a sustained-attention context (α = .05, power = .95, d′ = 0.80), yielded a required sample size of 23. However, due to scheduling, data collection yielded 24 usable participants. Results reported in this manuscript are based on the full sample (N = 24).

Fourteen participants (7 women and 7 men; *M*_age_ = 20.86 years, *SD*_age_ = 4.45) completed the EEG experiment. Instead of pre-determining required sample size, we used optional stopping rules and stopped data collection when Bayes Factor for the main effect of attention validity on SSVEPs exceeds 3 (Figure 4B; Rouder, 2014).

## Materials and Methods: Behavioral Study

### Apparatus

In a dark room, subjects viewed stimuli presented on a 27-in. IPS monitor (ASUS ROG VG279QM, 1920 × 1080 pixels at 144 Hz), located 63 cm in front of the participants (44.12 pixels per degree visual angle (dva). Stimuli were presented on a gray (42.5 cd/m^2^) background. Gaze position was recorded using EyeLink1000 desk-mounted tracker at a 1000 Hz sample rate. A nine-point grid was used for calibration before experiment, with the calibration area reduced to 45% of full screen to improve the precision near fixation. Auditory stimuli were delivered through two speakers (Creative SBS 4.1 450 system) placed immediately to the left and right of the monitor, distanced approximately 25 dva away from the center of the display. We used a chinrest and forehead rest to minimize participants’ head movement. The experiment display and eye-tracker were controlled using Psychophysics Toolbox (Brainard & Vision, 1997) and Eyelink toolbox (Cornelissen et al., 2002) in MATLAB.

### Procedure

#### Main experiment procedure

Each block consisted of a sequence of intermixed auditory and visual task trials (Figure 1A). For the auditory task, participants performed a dichotic listening task in which two simultaneous digit streams were played from speakers positioned to the left and right of the monitor. On each trial, one digit was spoken in a male voice and the other in a female voice, with the side assigned to each voice randomly alternating across trials. Before each block, participants viewed a written instruction to attend to digits from (1) the left speaker (attend-left), (2) the right speaker (attend-right), or (3) a specific voice regardless of speaker location (male or female; attend-voice). Within the attended stream, 20% of digits were immediate repetitions of the preceding digit. Participants were instructed to press the spacebar whenever they detected such a repetition (auditory 1-back task) while maintaining fixation on a central cross. For example, in the attend-left condition, participants ignored the right speaker and responded whenever the same digit appeared twice in a row in the left stream, regardless of voice. In the attend-voice condition, participants monitored digits spoken in the target voice across both speakers. Critically, this condition did not require spatial attention to either location, because the male and female voices were randomly distributed across both speakers within each block. This design allowed us to isolate the effect of auditory spatial attention while holding the sensory input constant across conditions. Spacebar responses were monitored for 1,200 ms following sound onset; the next digit played on schedule after a variable inter-stimulus interval (0–200 ms) regardless of whether participants had responded.

Within each block, participants also performed an intermittent visual orientation-discrimination task (Figure 1B), designed to probe visual processing across hemifields under different auditory attention conditions. On each visual trial, a near-threshold Gabor patch was briefly presented either to the left or right of fixation for 83.33 ms (twelve frames). Participants were instructed to report whether the Gabor was tilted counterclockwise or clockwise using the left and right arrow keys. Upon Gabor offset, the fixation cross turned blue to signal that a visual stimulus had appeared and to prompt a response. The visual task was self-paced: the experiment did not advance until participants made a response for visual trials, and they were instructed to guess if they had not perceived the Gabor patch.

Each block contained 156 auditory and 52 visual task trials (208 total), presented in a random order subject to two constraints: (1) each block began with at least five auditory trials before the first visual trial, and (2) at least two auditory trials separated any two consecutive visual trials. Across the main experiment, each of the three auditory attention conditions (attend-left, attend-right, and attend-voice) was completed three times. Of the three attend-voice blocks, at least one was assigned to each voice gender (male, female). Prior to the main experiment, participants completed four practice blocks, one for each attention condition (attend-left, attend-right, attend-male voice, attend-female voice), each containing 32 trials (24 auditory and 8 visual).

### Stimuli

#### Auditory Stimuli

Audio files of male and female voices articulating the ten digits (0–9) were generated using a freely available online AI text-to-speech converter. The resulting waveform audio files (.wav) were imported into MATLAB for preprocessing prior to the main experiment. Loudness was first normalized in accordance with the EBU R128 standard to achieve a target integrated loudness of −23 LUFS (Loudness Units Full Scale), using the *integratedLoudness.m* function. Subsequently, the speech segments were extracted from each audio file using the *detectSpeech.m* function (Table 1).

#### Visual Stimuli

A fixation cross was always presented at the center of the gray background (Figure 1B). Additionally, two outlined circles (placeholder), with 9.52 degrees of visual angle (dva) in diameter, were presented 22.52 dva to the left or right from the center so that they are located at the left- or right-most edge of the monitor. Note that both placeholders and Gabor patches were presented as peripheral as possible to maximize the spatial correspondence between visual stimulus and sound sources (two speakers), located right next to the monitor. In visual task trials, a tilted Gabor patch was presented within either left or right placeholder for the orientation discrimination task. The Gabor patch was generated by multiplying a sinusoidal grating of 0.8 cycles per degree with a Gaussian envelope with standard deviation of 1.7 dva. Each Gabor patch was tilted 20 degrees either counterclockwise or clockwise.

The contrast of the Gabor patch was individually determined through a contrast thresholding procedure before the main experiment. Subjects performed a block of the visual orientation discrimination task similar to the visual trials in the main experiment (Figure 1B). Each trial began with an initial delay period (500-1000 ms) with placeholders. Then, a Gabor patch was briefly presented either in the left or right visual hemifield and disappeared. Subjects reported whether the Gabor patch was tilted counterclockwise or clockwise. The Michelson contrast of the Gabor patch was titrated using an adaptive psychometric procedure to calculate the subthreshold contrast level. Specifically, we implemented the PEST (Pentland, 1980) method to approximate the contrast level required to achieve 75% orientation discrimination accuracy. The Palamedes toolbox (Prins and Kingdom, 2018) was used with a wrapper code (https://github.com/michaeljigo/palamedes_wrapper). Four adaptive titrations, each containing 24 trials, were randomly intermixed. A single subthreshold contrast level was obtained by averaging the contrast level used in last ten trials from each adaptive procedure. In case one of the adaptive procedures did not converge with visual inspection, we averaged contrast levels from the rest of the converged adaptive procedures.

### Analysis

#### Auditory task performance

For the auditory 1-back task, we defined *hit* as a ‘repeated’ (spacebar) response when the current digit in the attended stream was identical to the previous digit. In contrast, ‘repeated’ responses when the current digit was different from the last digit were defined as *false alarm*. The 1-back task performance was quantified as sensitivity (*d’*):

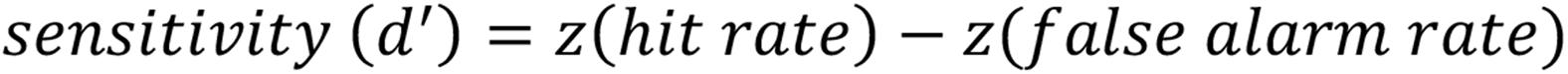

#### Visual task performance

Orientation discrimination performance was quantified as discrimination accuracy (chance level=0.5), computed separately for each Gabor patch location and auditory attention condition. To assess the effect of auditory spatial attention on visual discrimination accuracy, accuracy values were averaged according to the congruency between the attended auditory location and the location of the Gabor patch (Figure 2A). Trials in which the attended auditory location matched the Gabor patch location were labeled as valid trials, whereas trials in which the two locations differed were labeled as invalid trials (Figure 2B). In addition, discrimination accuracy in attend-voice blocks was calculated as a baseline to examine whether the observed effects of auditory spatial attention reflected facilitation or suppression. The same analyses were performed on response times measured on correct trials.

To further examine whether temporary shifts of attention following auditory repetitions contributed to subsequent changes in visual discrimination performance, trials were sorted according to the type of the immediately preceding auditory trial: (1) trials in which the preceding trial was a *repeated* auditory task trial, and (2) trials in which the preceding trial was a *non-repeated* auditory trial. Furthermore, to account for unequal number of trials sorted as repeated or non-repeated trials, we iteratively subsampled the non-repeated trials in a random manner so that the two conditions each contributed the same number of observations (Supplement Figure 1). After iterating subsampling 5,000 times, we obtained a distribution of t-statistics associated with the effect of attention validity on visual accuracy for non-repeated trials.

#### Eye tracking data

Raw EyeLink Data Format (EDF) files were analyzed to examine gaze shifts under different auditory spatial attention condition. Blinks were identified using the EyeLink parsing algorithm, and gaze samples within ±100 ms of each blink were discarded and linearly interpolated. Then, we extracted gaze position data during the pre-stimulus interval (−500 to 0 ms relative to visual stimulus onset). Distribution of horizontal gaze position (x) were fitted with normal distributions, and the means were used to compare the center of gaze. Positive and negative gaze shifts respectively indicated rightward and leftward shifts. We also extracted gaze position data across entire blocks and the post-visual-stimulus period (0 to 500 ms relative to visual stimulus onset) and performed the same analyses (Supplementary Figure 2).

## Materials and Methods: EEG Study

### Apparatus

All visual stimuli were presented on a ViewPixx monitor (1,920 × 1,080 pixels at 120 Hz; VPixx Technologies) placed 38 cm from the participant (39.10 pixels per degree visual angle) in a dark room. Stimuli were presented on a gray (43.3 cd/m^2^) background. Auditory stimuli were delivered through two speakers (Creative SBS 4.1 450 system) placed immediately to the left and right of the monitor. The experiment display and eye-tracker were controlled using Psychophysics Toolbox (Brainard & Vision, 1997) and Eyelink toolbox (Cornelissen et al., 2002) in MATLAB.

Scalp electric activity was recorded with 32-channel Ag/AgCl electrode cap (Brain Products, GmbH) connected to BrainVision actiChamp amplifier. Signals were referenced to the right mastoid online, band-pass filtered online between 0.01 to 112.5 Hz, and digitized at 500 Hz. To improve sampling of posterior neural activity, we used custom electrode configuration modified from the 10-20 system by re-locating three frontal electrodes (FC5, FCz, FC6) to the occipital scalp (I3, Iz, I4 respectively). Horizontal eye movements were monitored with a bipolar HEOG electrode pair placed at the outer canthi of each eye; an additional reference electrode on the right side of the neck served as ground for the HEOG signal. Impedances were kept below 20 kΩ for all channels before recording began.

### Procedure

Participants performed an auditory 1-back task in a dichotic listening setup while passively viewing two flickering checkerboard patterns presented in the left and right visual hemifield (Figure 3A). During each 10-second trial, two independent streams of digits were played simultaneously from speakers positioned to the left and right of the monitor. Each digit was spoken in either a male or female voice, selected randomly. Before each trial, participants viewed a written cue instructing them to attend to either the left auditory stream (attend-left) or the right auditory stream (attend-right). The trial began when participants pressed the arrow key corresponding to the cued side (left or right arrow). Each trial contained eight digits per stream, with an inter-stimulus interval of 0.9–1.1 seconds and a 20% repetition rate. Participants were instructed to press the spacebar whenever an immediate repetition occurred in the attended stream. Concurrent to the auditory task, two circular checkerboard stimuli, positioned to the left and right of fixation, flickered at either 12 Hz or 15 Hz frequency. These visual stimuli were entirely task-irrelevant and thus participants were instructed to ignore them and maintain fixation at the center of the display.

Depending on the location of the attended auditory stream and the assignment of two frequencies tagged onto each visual hemifield, there were four different types of trials (Figure 3B). Different combinations differed on which flickering rate of visual stimuli was either valid or invalid to the auditory spatial location. For example, if the checkerboard pattern on the left visual field was flickering at the rate of 12 Hz (right stimuli flickering at the rate of 15 Hz), 12 Hz was labeled as valid frequency when subjects were attending to left auditory stream, and invalid when attending to the right auditory stream. Each trial type was repeated 50 times, resulting in 200 trials randomly presented across the experiment.

### Stimuli

#### Auditory Stimuli

Audio stimuli for the dichotic listening task were identical to that used for behavioral experiment.

#### Visual Stimuli

Two radial checkerboard patterns were positioned to the left and right of the central fixation cross (Figure 3C, top panel). The center of each stimulus was 16.55 dva lateral from fixation. Each checkerboard, subtended 8° of visual angle in diameter, was comprised of 5 concentric radial steps and 8 angular wedges (each wedge spanning 22.5°). The two patterns flickered by alternating black–white contrast at either 12 Hz or 15 Hz. The flicker frequency assigned to the left and right stimulus was counterbalanced across blocks, ensuring that each hemifield was equally often associated with each frequency. The actual flicker rate of each stimulus was verified with a photodiode placed over the screen, allowing us to confirm that the delivered frequencies matched the intended 12 Hz and 15 Hz values. A central fixation cross remained on screen throughout each trial, and participants were instructed to maintain fixation for its entire duration.

### Analysis

EEG data were preprocessed using the EEGLAB toolbox (Delorme & Makeig, 2004) and ERPLAB toolbox (Lopez-Calderon & Luck, 2014) in MATLAB (The MathWorks, Natick, MA, USA). Signals from noisy electrodes were interpolated using the *eeg_interp.m* function^3^. Next, we performed Independent Component Analysis (ICA) to remove EEG artifacts associated with eye blinks and eye movements (Delorme & Makeig, 2004; Drisdelle et al., 2017). On average, 2.43 components were identified and removed per participant. A detailed description of the removed components for each participant can be found on OSF. The preprocessed EEG data were re-referenced to the average across all electrodes. Lastly, EEG data were epoched for each trial from −500 to 10,000 ms relative to trial onset and filtered with a high-pass (> 0.01 Hz) and a low-pass (< 80 Hz) filters. Epochs were baseline-corrected using the mean signal from −200 to 0 ms relative to trial onset.

For SSVEP analysis, we applied a Fourier transform to neural activity averaged across trials sharing the same attended location and tagged-frequency location (500–9,500 ms per trial; Figure 3C). SSVEP amplitude was defined as the absolute value of the complex Fourier coefficient, extracted as the maximum amplitude within ±0.2 Hz of each stimulation frequency (12 Hz and 15 Hz).

For each stimulation frequency and hemifield assignment, SSVEP amplitudes were normalized across the two attention conditions (attend-left, attend-right) by subtracting and then dividing by their mean, yielding a relative measure of how much attention modulated the response at each frequency, independent of baseline differences in absolute amplitude across frequencies or hemifields (Andersen et al., 2011; Störmer et al., 2014):

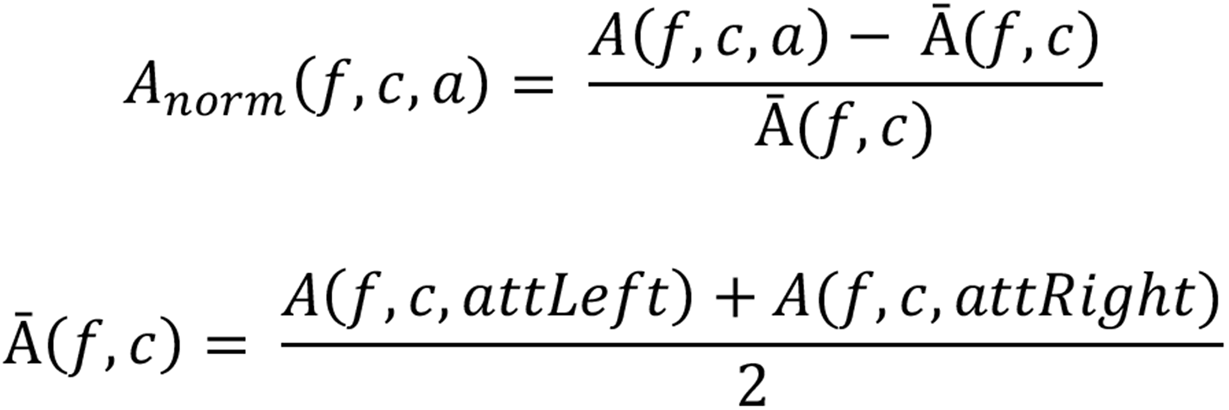

where *A*(*f*, *c*, *a*) is the amplitude at frequency *f*, under hemifield assignment *c*, when attention was directed to location *a*. This yielded four independent normalization groups, each normalized separately across its own attend-left/attend-right pair. Normalized amplitudes were then labeled as valid when the frequency’s hemifield matched the attended location and invalid otherwise, then averaged across a set of occipital electrodes (I3, Iz, I4, O1, Oz, O2, PO7, PO3, POz, PO4, PO8) for statistical comparison (Störmer et al., 2014; Figure 4C).

We computed posterior alpha-band activity over electrodes contralateral and ipsilateral to the attended auditory location (Feng et al., 2017; Keefe & Störmer, 2021; Störmer et al., 2016). Unlike SSVEP responses, which are phase-locked to the flickering rate of the visual stimuli and therefore computed by applying a Fourier transform to trial-averaged EEG data, endogenous alpha oscillations are not phase-locked across trials. Accordingly, Fourier transforms were applied to single-trial EEG data to estimate alpha-band amplitude.

We focused on parietal–occipital electrodes (PO7, PO8, P7, P8; Keefe & Störmer, 2021) and defined the alpha band as 8–11.5 Hz to avoid spectral overlap with the tagging frequencies (12 and 15 Hz). Alpha-band amplitudes were averaged within this range and submitted to a lateralization index (LI):

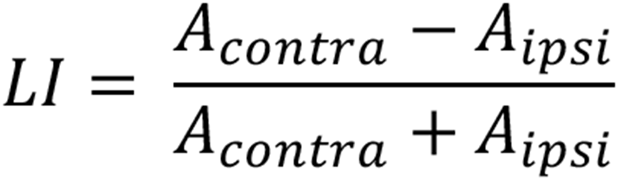

Where *A_contra_* and *A_ipsi_* denote the average alpha-band amplitude when attending to contralateral and ipsilateral hemifield, respectively. A negative LI therefore reflects reduced alpha amplitude over electrodes contralateral to the attended hemifield. For both statistical analyses and topographical mapping, amplitudes were collapsed across left-and right-hemisphere electrode pairs.

## Acknowledgments and funding sources

This work was supported by the National Science Foundation (NSF BCS-2446115).

## Author Contributions

Y.M.C. and V.S.S. designed the research; V.S.S. supervised the research; Y.M.C. performed the research, collected and analyzed the data; Y.M.C. and V.S.S. wrote the paper.

---

**Supplementary Figure 1.**
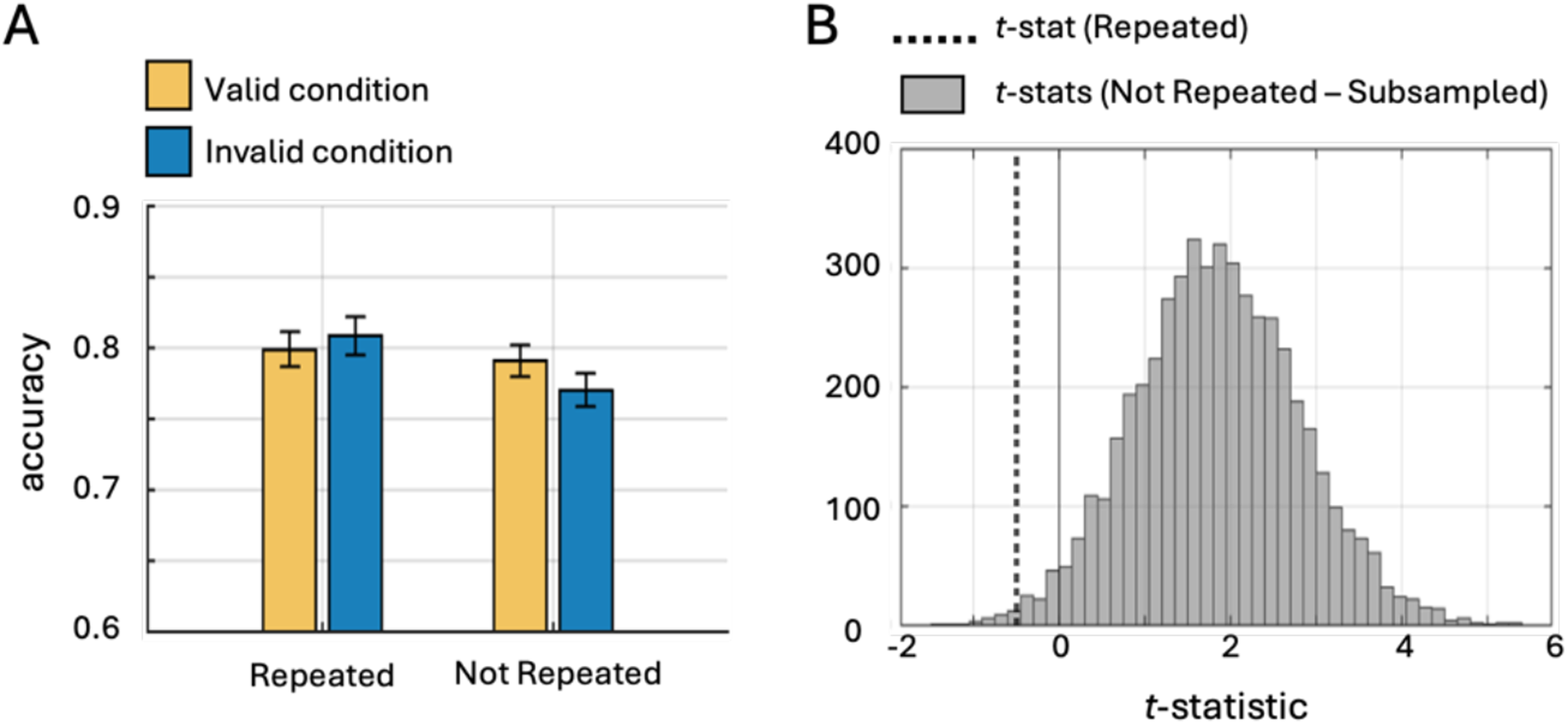
Additional analysis to account for temporary shifts of attention. To test an alternative account of cross-modal facilitation — namely, that temporary shifts of auditory attention following repeated trials drive the effect — we sorted visual trials according to the nature of the immediately preceding auditory event: repeated (the 1-back target) or not-repeated. This analysis revealed a significant effect of auditory spatial attention only in not-repeated trials, but not in repeated trials (Figure 2C). However, because only 20% of auditory digits were repeated, the repeated condition contained far fewer trials (M = 61.9, SD = 5.9) than the not-repeated condition (M = 250.1, SD = 5.9), which could have led to asymmetric statistical power. To account for this imbalance, we randomly subsampled the not-repeated trials to match the number of repeated trials. (A) In one representative subsample, the interaction between attention validity and preceding-trial type remained significant (F(1, 23) = 5.48, p = 0.028, *η_p_^2^* = 0.19), reproducing the pattern obtained without subsampling. (B) We repeated the subsampling 5,000 times and computed paired-samples t-statistics contrasting valid and invalid cues for each iteration. The resulting distribution of t-statistics was strongly shifted above zero, with 97.4% of iterations yielding a positive t-value, and only 0.6% producing a t-value smaller than that observed in the repeated-trial condition (t = −0.498). This bootstrapped result confirms that cross-modal facilitation cannot be attributed to brief temporary shifts of attention, and supports the conclusion that sustained, goal-directed auditory spatial attention systematically enhances visual processing at the attended location.

**Supplementary Figure 2.**
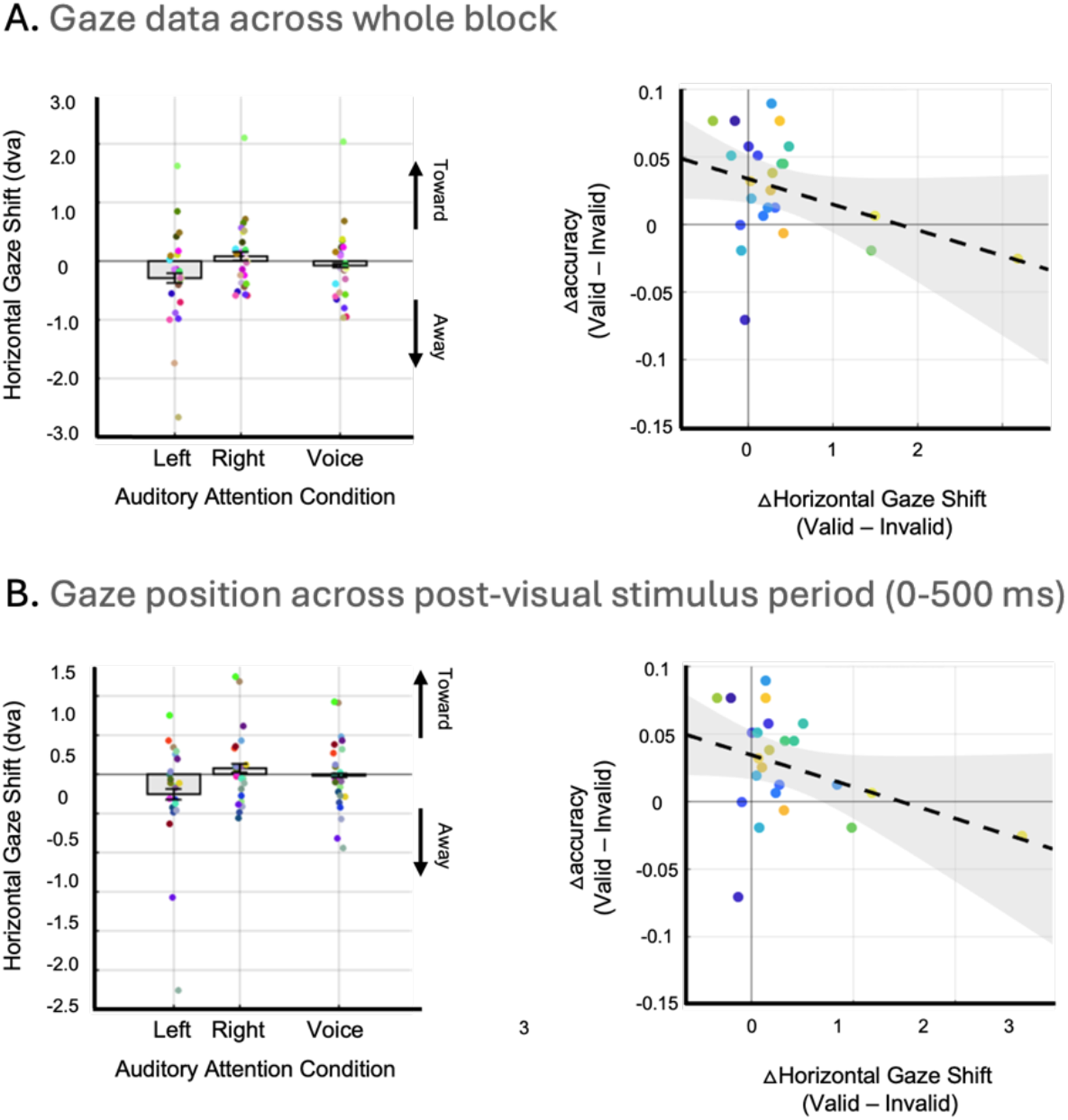
Gaze position analysis using alternative time windows. Identical analyses were performed using gaze position data averaged across either the entire block (A) or the post-visual-stimulus period (B; 0–500 ms), yielding the same pattern of results as the pre-stimulus analysis reported in the main text. For each time window, the left panel shows horizontal gaze shifts, with negative and positive values indicating shifts toward the left and right visual field, respectively, and the right panel shows the relationship between gaze shift magnitude and the effect of sustained auditory attention on visual discrimination performance. Each dot represents one participant; error bars indicate within-subject standard error.

**Supplementary Figure 3.**
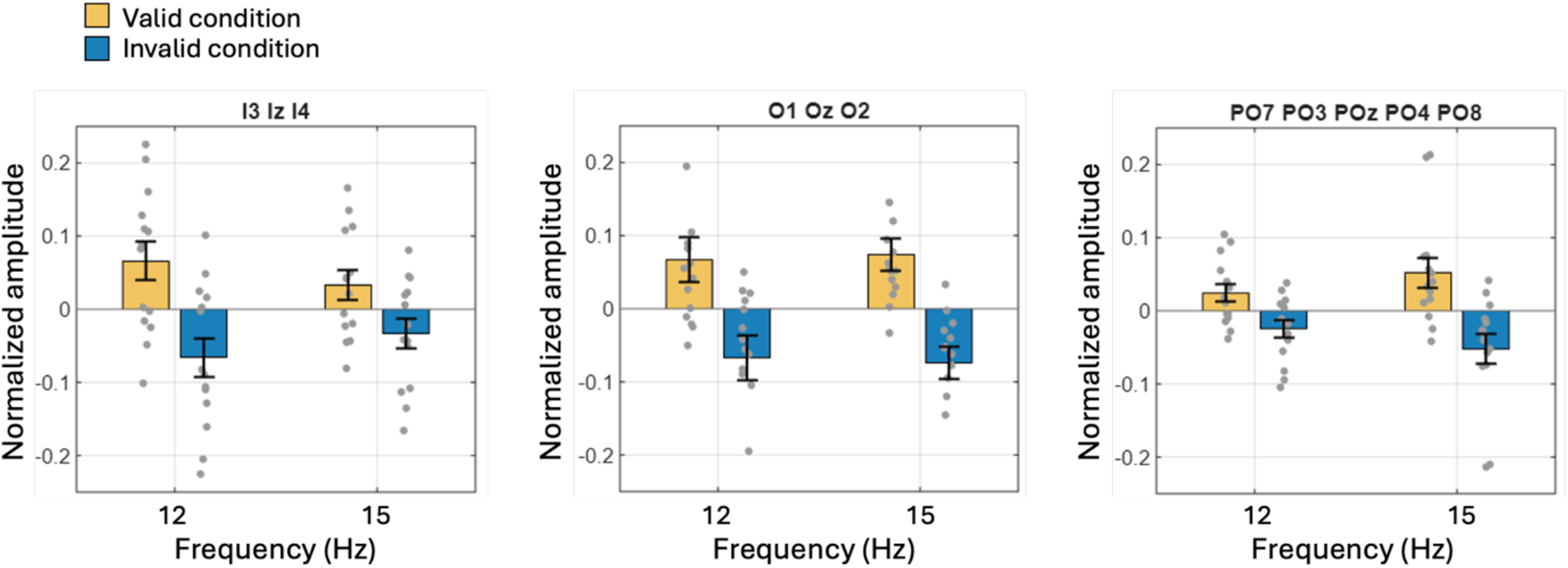
SSVEP amplitude results per electrode group. Normalized SSVEP amplitude for each electrode group: (A) I3, Iz, I4; (B) O1, Oz, O2; (C) PO7, PO3, POz, PO4, PO8. A 2×2×3 repeated-measures ANOVA revealed a significant main effect of attention validity (*F*(1, 13) = 11.64, *p* = .005, *η_p_^2^* = 0.472, BF_incl_ = 10.63), with no significant attention validity × electrode group interaction (*F*(2, 26) = 1.205, *p* = .316, *η_p_^2^* = 0.085, BF_incl_ = 0.592).

## Footnotes

1 We also performed the same analyses using gaze position data averaged across the entire block or the post-visual-stimulus period (0–500 ms), and found the same pattern of results in both cases (Supplementary Figure 2).

2 Because normalization was applied separately within each frequency condition, the mean normalized amplitude was zero by construction for both conditions; the main effect of tagged frequency is therefore uninterpretable and is not reported.

3 Interpolated channels for each subject: I3 for subject 1; I3 for subject 6; C3 for subject 9; CP1 for subject 12; CP3, CP5 for subject 13; M1 for subject 14

